# Cost-effective hybrid long- and short-read sequencing enables accurate somatic structural variant detection

**DOI:** 10.64898/2026.02.16.706063

**Authors:** Runtian Gao, Tao Jiang, Heng Hu, Zhongjun Jiang, Shuqi Cao, Murong Zhou, Yuming Zhao, Guohua Wang

**Affiliations:** College of Life Science, Northeast Forestry University, Harbin 150000, China; College of Computer and Control Engineering, Northeast Forestry University, Harbin 150000, China; Faculty of Computing, Harbin Institute of Technology, Harbin 150001, China; Zhengzhou Research Institute, Harbin Institute of Technology, Zhengzhou 450000, China

## Abstract

Somatic structural variant (SSV) calling typically requires matched normal data. Incorporating relatively inexpensive short-read sequencing not only provides essential germline information but can also replace a substantial portion of long-read sequencing, thereby enabling more cost-effective somatic SV detection. Here, we present SomaSV, a hybrid sequencing framework that integrates 30× tumor long-read data with matched normal data comprising 10× long-read and 30× short-read sequencing. This design achieves high-accuracy somatic SV detection while remaining cost-competitive. Comprehensive benchmarking demonstrates that SomaSV outperforms current state-of-the-art methods by more than 13% in F1 score while reducing sequencing costs by approximately 19%. Moreover, SomaSV identifies clinically relevant cancer-associated genes, including *CLDN4* and *ROBO2*, highlighting its potential for discovering valuable biomarkers to support early cancer screening and diagnosis. The source code for SomaSV can be accessed at https://github.com/eioyuou/SomaSV.

## Introduction

Somatic structural variants (SSVs) are genomic rearrangements acquired during tumorigenesis and evolution, existing in either clonal or subclonal forms [1]. These include deletions, insertions, inversions, duplications, and complex combinations thereof. SSVs can drive tumor progression by altering gene dosage, disrupting coding sequences, or reshaping cis-regulatory elements, thereby influencing tumor classification, prognostic assessment, and therapeutic target identification [2]. Long-read sequencing (LRS) has substantially improved the resolution of complex structural variants, enabling haplotype-aware reconstruction of rearrangement events in cancer genomes [3]. Accordingly, several LRS-based SSV detection tools, such as Severus [4], nanomonsv [5], SAVANA [6], and SVision-pro [7], achieve robust performance under high-coverage LRS data. Incorporating relatively inexpensive short-read sequencing (SRS) can provide essential germline information and partially substitute for long-read data, offering the potential for more cost-effective somatic SV detection [8]. Despite this potential, an effective framework that achieves this balance has not yet been established.

In this study, we present SomaSV, a hybrid framework for detecting SSVs using tumor LRS and matched normal LRS and SRS data (Fig. 1a). SomaSV comprises two key modules: an LRS-only mode, which analyzes tumor and normal LRS data, and a standard mode, which incorporates matched SRS from the normal sample to improve germline discrimination.

**Fig. 1.**
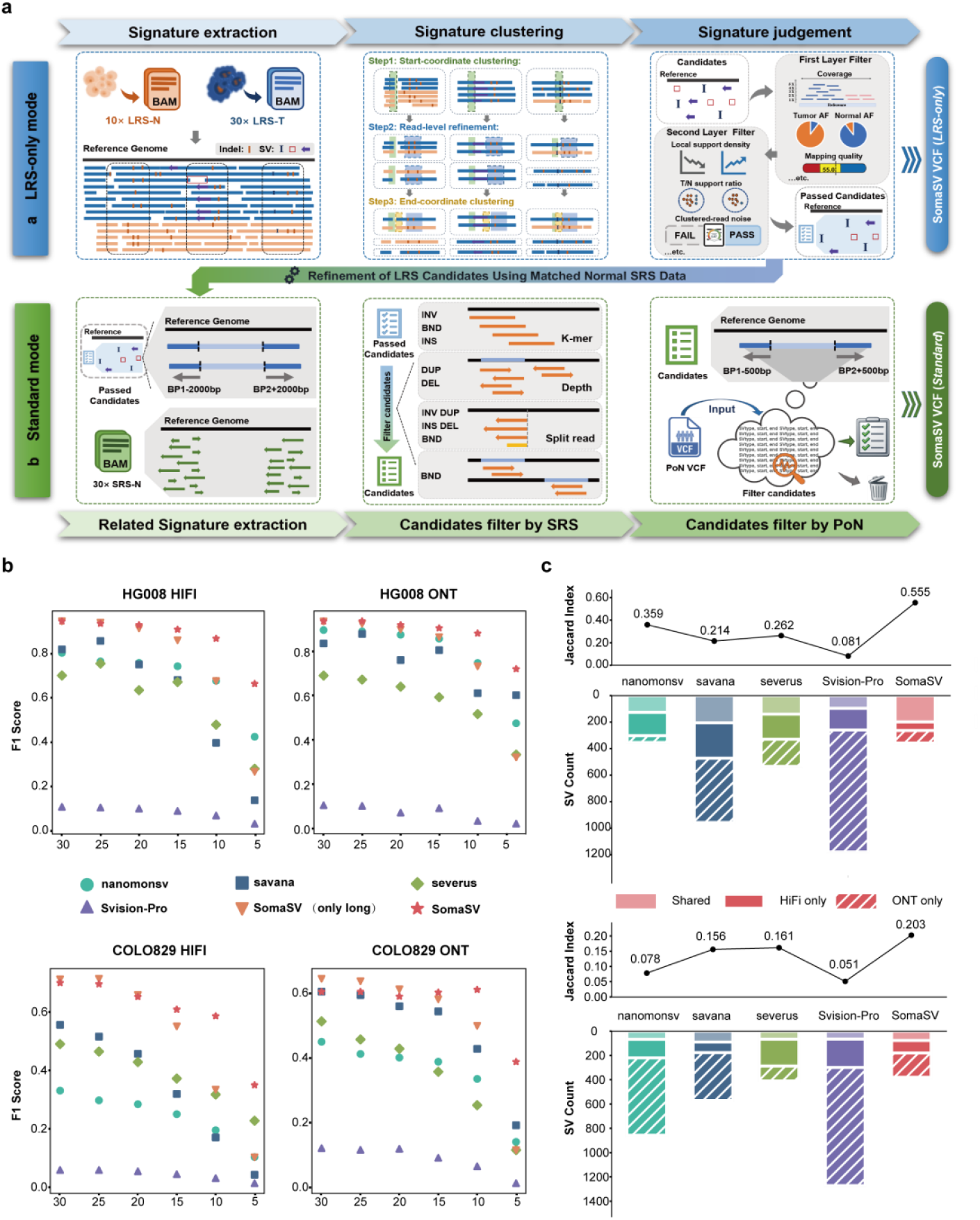
SomaSV workflow and benchmarking across normal LRS depths and sequencing platforms. **a**. Schematic of the SomaSV framework, comprising an LRS-only module for candidate discovery and a standard module that incorporates normal SRS and PoN filtering for somatic SV refinement. **b**. F1 scores for HG008 and COLO829 under 30× tumor LRS and varying normal LRS (30× to 5×), with or without 30× normal SRS augmentation. **c**. Cross-platform consistency between HiFi and ONT callsets at 30× tumor and 10× normal LRS, measured by Jaccard index and SV count.

In the LRS-only mode, SomaSV extracts read-level breakpoint signals from tumor and normal LRS data and clusters them into candidate SV loci using a multi-stage strategy. Candidates are filtered using a variant allele frequency-aware (VAF-aware) approach with adaptive thresholds on coverage, read support, and tumor–normal imbalance. A feature-based model then scores filtered variants by integrating read- and breakpoint-level metrics, and high-confidence SVs are retained based on a predefined threshold.

In the standard mode, matched normal SRS data are used to refine LRS candidates through orthogonal validation, incorporating features such as k-mer concordance, depth profiling, and breakpoint-aligned short-read support within ±2 kb of predicted breakpoints. This additional SRS-derived evidence improves resolution and reduces false positives, particularly in settings with low-depth normal LRS or cost constraints. Notably, both modes optionally support panel-of-normals (PoN) filtering, which removes recurrent artifacts and rare germline variants based on breakpoint proximity.

## Results and Discussion

To establish a consistent benchmark, we evaluated all methods using 30× long-read sequencing data for both tumor and normal samples. Analyses were performed on two tumor cell lines (HG008-T from a pancreatic ductal adenocarcinoma, COLO829 from melanoma fibroblasts) and their matched pair (HG008-N-P from normal pancreatic, COLO829BL from normal B lymphoblasts) [9,11]. We present HG008 as the primary case study, with COLO829 used to confirm trend consistency. Under this setting, SomaSV achieved the highest F1 scores: 94.37% (HiFi) and 91.37% (ONT) for HG008, and 72.83% (HiFi) and 65.13% (ONT) for COLO829 (Fig. 1b). This superior performance reflects SomaSV’s design, which incorporates VAF-aware filtering and multi-feature scoring to prioritize true somatic events, even under conditions where high LRS depth benefits all methods.

While high tumor coverage is often necessary to overcome heterogeneity and low VAF, the effect of normal coverage on SSV detection is less clear. We therefore fixed tumor LRS at 30× and downsampled normal LRS from 30× to 5×. we fixed tumor LRS at 30× and systematically reduced normal LRS from 30× to 5× to assess its impact on detection performance. As shown in Fig. 1b, most tools exhibited a progressive decline in F1 score as normal coverage decreased, often accompanied by a substantial increase in false positives. SomaSV maintained high accuracy (F1: 94.37%–85.70% for HG008, 72.83%-56.30% for COLO829) between 30× and 15× normal coverage, despite reduced depth. This relative robustness under coverage imbalance is attributable to its algorithmic design, which incorporates VAF–stratified filtering and feature-level scoring to prioritize high-confidence somatic variants. Performance declined more markedly at 10× and 5×, indicating that additionally essential germline mutation signals need to be involved in order to discriminate SSVs accurately. It seems that this situation can be eased through high-coverage but low-cost SRS data for a normal sample.

To evaluate whether SRS can compensate for limited normal LRS coverage, we systematically assessed the effect of incorporating SRS data from the normal sample across a range of downsampled conditions. When both tumor and normal samples were sequenced at 30× LRS, the addition of 30× SRS from the normal sample had negligible effect on detection accuracy. In contrast, as the normal LRS depth decreased, the benefit of SRS became increasingly evident (Fig. 1b). At 10× normal HiFi coverage, supplementing with 30× SRS improved the F1 score of SomaSV from 67.18% to 86.49%, primarily by increasing precision from 52.11% to 81.78% while maintaining high recall. This improvement represented the largest gain observed across all hybrid configurations. However, when normal long-read coverage was further reduced to 5×, overall performance degraded substantially despite the inclusion of SRS. Under this setting, SomaSV achieved only 65.93% F1 and 53.60% precision, and all comparator tools exhibited precision values below 30%, suggesting that SRS alone cannot fully rescue performance under severely depleted long-read conditions. These findings support a hybrid configuration of 30× tumor LRS, 10× normal LRS, and 30× normal SRS as a cost-effective strategy that maintains high accuracy while substantially reducing long-read sequencing demands on the normal sample.

Given that long-read sequencing platforms differ in their error profiles, which can introduce platform-specific artifacts and impact somatic SV detection, we next evaluated cross-platform robustness by comparing callsets derived from HiFi and ONT sequencing data of the sample. To ensure comparability across conditions, we adopted a standardized configuration of 30× tumor LRS, 10× normal LRS and 30× normal SRS for both platforms in all subsequent analyses. We quantified cross-platform concordance using the Jaccard index [10], defined as the ratio of the intersection to the union of two callsets, which reflects global consistency in SV prediction. Among all evaluated methods, SomaSV achieved the highest cross-platform concordance in HG008, with a Jaccard index of 55.5%, substantially outperforming nanomonsv (35.9%), Severus (26.2%), and SVision-Pro (8.1%) (Fig. 1c). Similar trends were observed in COLO829, where SomaSV again yielded the highest overlap between platforms. Since short reads are unaffected by long-read-specific error modes, their inclusion helps correct platform-induced artifacts. These results highlight the utility of SRS in improving cross-platform consistency in somatic SV detection.

To evaluate the false positive tolerance of each method, we generated synthetic tumor–normal pairs by randomly partitioning the original normal LRS data into two disjoint read sets, treated as “tumor” and “normal.” As both originate from the same genome, any detected somatic SVs reflect noise, mapping errors, or algorithmic artifacts. Across four datasets and two platforms, SomaSV consistently reported the fewest false positives, while other tools often produced substantially more spurious calls under identical conditions (Fig. 2a). These results indicate that methods relying solely on long-read data are more prone to platform-specific noise, such as alignment errors and indels. By incorporating orthogonal SRS signals, SomaSV effectively suppresses such artifacts. In addition to improving specificity, short-read integration helps correct for systematic biases across platforms, enhancing robustness in low-signal or error-prone settings.

**Fig. 2.**
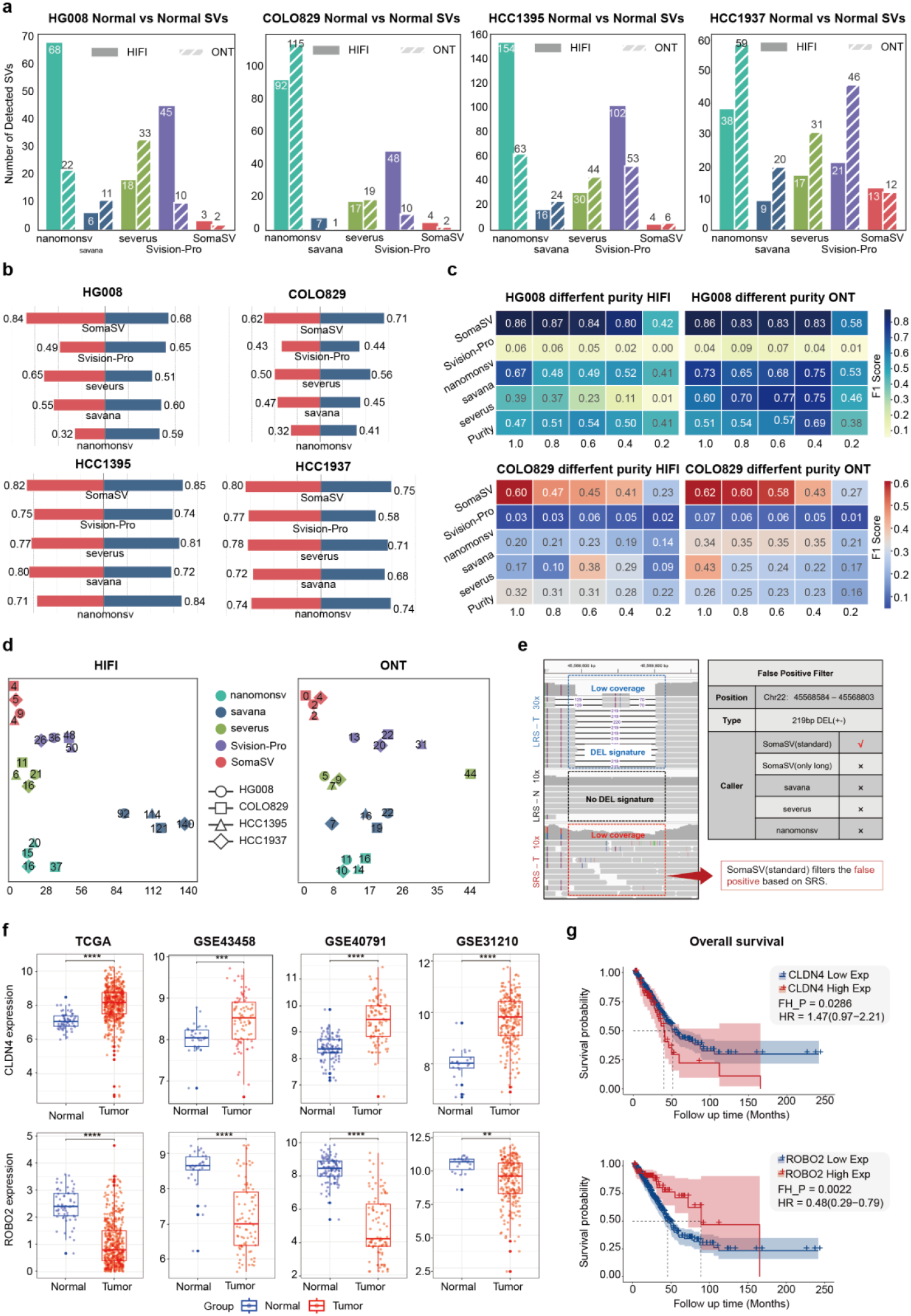
Robustness, specificity, and functional relevance of SomaSV. **a**. False positives in normal–normal comparisons across four samples (HG008, COLO829, HCC1395, HCC1937) oval of artifactual calls by SomaSV through short-read–based filtering in the standard mode. **b**. Reproducibility assessed by Jaccard overlap between somatic SV callsets from two independent 30× tumor down samplings. **c**. F1 scores across tumor purity series (0.8–0.2) show SomaSV’s robustness under subclonal conditions in HG008 and COLO829 (HiFi/ONT). **d**. Proxy analysis of germline leakage: overlap between somatic SVs and high-confidence germline SV sets. **e**. Representative locus showing removal of artifactual calls by SomaSV through short-read–based filtering in the standard mode. **f**. Tumor–normal differential expression of *CLDN4* and *ROBO2* across TCGA and GEO cohorts. **g**. Kaplan–Meier survival analysis stratified by *CLDN4* and *ROBO2* expression shows association with patient prognosis.

To assess the robustness of somatic SV calling under stochastic variation in sequencing data, we evaluated replicate concordance across independently downsampled tumor datasets. For each sample, two distinct 30× subsets were randomly drawn from the original tumor BAM and analyzed against the same matched 10× normal LRS and 30× normal SRS. The consistency of SV predictions between replicates was quantified using the Jaccard index (Fig. 2b). Across all four datasets, SomaSV achieved the highest concordance under both HiFi and ONT platforms, including Jaccard scores of 84.35% and 68.13% in HG008. In contrast, other tools showed more variable performance. These results demonstrate that SomaSV maintains high reproducibility under random sampling variation, in part due to the fine-scale resolution provided by short-read evidence, which helps anchor SV breakpoints and mitigate errors propagated from long-read alignment uncertainty.

In addition to normal coverage constraints, tumor purity poses another major challenge to somatic SV detection. To evaluate the sensitivity of SSV detection under realistic low-signal conditions, we simulated tumor samples at varying purities by mixing 60× tumor and 30× normal LRS to a fixed 30×, and analyzed each against a matched 10× normal LRS and 30× normal SRS. This setup reflects a challenging scenario where both tumor purity and control coverage are limited, leading to reduced signal contrast and elevated noise. SomaSV consistently outperformed other methods across all purity levels and platforms, with particularly notable advantages at 20% purity, where F1 scores remained at 0.42 (HiFi) and 0.58 (ONT) in HG008, while other tools failed to recover sufficient signal (Fig. 2c). Importantly, those additional information from SRS improve breakpoint resolution and germline SV delineation in the control, thereby sharpening tumor–normal contrast and enabling reliable somatic SV detection even under combined purity and coverage constraints.

For some samples without validated SSV truth sets, two tumor cell lines (HCC1395 from breast invasive ductal carcinoma, HCC1937 from breast invasive ductal carcinoma) and their matched normal pairs (HCC1395BL and HCC1937BL from peripheral blood lymphocytes) [11], we adopted an indirect but conservative evaluation strategy (Fig. 2d). High-confidence germline SV sets were generated by integrating calls from four germline SV tools on the matched normal. We then quantified the number of overlaps between each somatic callset and the germline SV set, using this as a proxy for misclassification risk. Across both HiFi and ONT, SomaSV consistently showed the lowest overlap, indicating better suppression of germline signals. This reflects the benefit of SRS-informed germline filtering, which helps eliminate false somatic calls arising from ambiguous or under-resolved germline SVs.

Finally, we applied SomaSV to a lung adenocarcinoma sample (H2009) [11] to evaluate its ability to detect clinically relevant somatic SVs in a real tumor context. Comparative analysis revealed multiple high-confidence variants uniquely identified by SomaSV, including a duplication of the oncogene *CLDN4* [12] and a deletion of the tumor suppressor *ROBO2* [13], which were not detected by other state-of-the-art somatic SV callers. Both genes exhibited consistent expression dysregulation across TCGA-LUAD and three independent GEO cohorts, with *CLDN4* significantly upregulated and *ROBO2* downregulated in tumors compared to normal tissues (Fig. 2f). Moreover, their expression levels were significantly associated with patient survival (Fig. 2g), implicating potential clinical relevance. These SVs likely contribute to tumorigenesis by altering gene dosage or disrupting regulatory domains. By enabling the discovery of functionally and clinically important SVs that are missed by existing methods, SomaSV provides a robust and sensitive framework for somatic genome interpretation.

In summary, SomaSV is an accurate SSV detection approach from cost-effective hybrid long- and short-read sequencing. Incorporating SRS data from the matched normal sample improves somatic specificity, helps suppress recurrent artifacts, enhances robustness across coverage and purity gradients, and increases cross-platform. Moreover, the inclusion of a PoN (gnomadSV v4.1) [14] further increases precision by filtering rare germline events and recurrent platform-specific noise. SomaSV also reduces the dependence on high-depth long-read sequencing in the normal sample, offering a cost-effective yet accurate strategy for somatic SV discovery. Future work would develop subclone-aware SSV calling through the integration of additional sequencing signals.

## Methods

### Performance benchmarking methodology

#### SSV detection benchmark in HG008 ground truth

The tumor–normal long-read and short-read datasets for HG008 were obtained from the Genome in a Bottle (GIAB) Consortium, hosted by the National Institute of Standards and Technology (NIST), as part of a project dedicated to the comprehensive characterization of benchmark cancer genomes. Matched tumor and normal samples were sequenced using PacBio HiFi, ONT, and Illumina short-read platforms. The associated benchmark somatic variant set was curated by GIAB and used as a reference for performance evaluation (NIST_HG008-T_somatic-stvar_DraftBenchmark_V0.3-20250220).

#### SSV detection benchmark in COLO829 ground truth

For the COLO829 cell line, Illumina and Oxford Nanopore sequencing data were obtained from the CASTLE project (https://github.com/CASTLE-Panel/castle), which provides somatic SV callsets and multi-platform whole-genome sequencing data from melanoma tumor and matched B lymphoblastoid normal samples. PacBio HiFi data for the same tumor–normal pair were retrieved from the PacBio public dataset repository (https://www.pacb.com/connect/datasets/) under the tumor/normal category. For evaluation, we used the updated consensus set of 54 somatic structural variants defined for COLO829/COLO829BL [15], which integrates results from four long-read replicates and includes both GRCh38- and CHM13-T2T-based coordinates.

#### Cross-platform consistency analysis

To evaluate cross-platform consistency in somatic structural variant detection, we analyzed callsets generated from PacBio HiFi and ONT sequencing data for the HG008 and COLO829 tumor–normal pairs. All samples were processed using a standardized configuration consisting of 30× tumor long-read sequencing, 10× normal long-read sequencing, and 30× normal short-read sequencing. Somatic SVs were independently called on each platform using SomaSV, nanomonsv, Severus, and Savana, with consistent parameters across platforms. For each tool, we used SURVIVOR (version 1.0.7) to perform pairwise merging of SV callsets from HiFi and ONT, allowing a maximum breakpoint distance of 500 base pairs and requiring matching SV types. Cross-platform concordance was measured using the Jaccard index, calculated as the number of shared SVs divided by the total number of unique SVs identified across both platforms.

#### False positive tolerance assessment

To evaluate the false positive tolerance of each method, we created synthetic tumor–normal pairs by equally splitting normal long-read sequencing data into two disjoint subsets using samtools (v1.19), assigning one to “tumor” and the other to “normal.” As both originate from the same genome, any detected somatic SVs represent false positives caused by sequencing noise, alignment artifacts, or algorithmic errors. This evaluation was conducted on HG008 and COLO829 using PacBio HiFi and ONT data, with 30× coverage applied to both pseudo-tumor and pseudo-normal samples.

#### Replicate concordance analysis

To assess the robustness of somatic SV calling to stochastic sampling variation, we evaluated replicate concordance using independently downsampled tumor datasets. For each sample, two distinct 30× tumor subsets were randomly drawn from the original long-read alignment file using samtools with different random seeds. Each replicate was analyzed against the same matched normal data, consisting of 10× long-read and 30× short-read coverage. Somatic SVs were called using SomaSV, nanomonsv, Severus, and Savana with consistent parameters across replicates. Concordance was quantified using the Jaccard index, calculated as the ratio of shared SVs to the total number of unique SVs across the two replicates. All sampling procedures and evaluation commands are described in Supplementary Note.

#### Tumor purity simulation and sensitivity evaluation

To evaluate the sensitivity of somatic SV detection under varying tumor purity, we created simulated tumor samples by mixing high-coverage tumor and normal long-read sequencing data. Specifically, 60× tumor and 30× normal PacBio HiFi or ONT reads were combined at predefined ratios to generate composite datasets with effective tumor purities of 100%, 80%, 60%, 40%, and 20%, each downsampled to a fixed 30× total coverage using samtools. Each simulated tumor dataset was analyzed against a matched 10× normal long-read and 30× normal short-read alignment. SV calling was performed using SomaSV, nanomonsv, Severus, and Savana under identical parameters across all conditions. Sensitivity was quantified using truvari (v5.0.3) by comparing each predicted callset to the reference somatic SV benchmark, with a 100 bp breakpoint tolerance and matching SV type required.

#### Germline overlap analysis in samples without somatic truth sets

For tumor–normal pairs without validated somatic SV truth sets, including HCC1395 and HCC1937 along with their matched normal lymphoblastoid controls HCC1395BL and HCC1937BL, we employed an indirect evaluation strategy based on overlap with high-confidence germline variants. Germline SVs were identified by applying four independent germline callers: Sniffles2, cuteSV, SVIM, and DeBreak, to the matched normal long-read data. Each tool was executed using its recommended default settings. The resulting callsets were merged using SURVIVOR, with the merging criteria requiring consistent SV types, a maximum breakpoint distance of 100 base pairs, and support from all four tools. The merged callset served as a proxy for germline structural variation. For each somatic SV detection method under evaluation, we quantified the number of predicted somatic SVs that overlapped with the germline proxy set. A lower number of overlaps was interpreted as stronger suppression of germline signal.

#### Functional and clinical relevance analysis

To evaluate the clinical and functional relevance of somatic SVs uniquely identified by SomaSV, we applied it to a primary lung adenocarcinoma sample (H2009) and compared the resulting SVs with those detected by three other state-of-the-art somatic SV callers. SVs uniquely identified by SomaSV were determined using SURVIVOR to merge callsets across tools, retaining only variants exclusive to SomaSV. These SVs were annotated using AnnotSV (v3.2.2) to identify affected genes. Expression analysis was then performed for selected genes across TCGA-LUAD and three independent lung adenocarcinoma datasets from GEO (GSE40791, GSE43458, and GSE31210). For TCGA-LUAD, RNA-seq and clinical data were downloaded from the UCSC Xena platform, and only primary tumor (01A) and control (11A) samples were retained, yielding 513 tumor and 58 normal samples, with 500 tumor samples having survival data. Differential expression analysis was conducted using DESeq2 (v1.49.9) with a significance threshold of |log2FC| > 0.5 and P < 0.05. For GEO datasets, gene expression matrices and group labels were processed using a unified R workflow, and differential expression was assessed with limma (v3.58.1) using the same thresholds. Prognostic relevance was evaluated using TCGA survival data. Patients were stratified into high- and low-expression groups based on optimal expression cutoffs for each gene, and Kaplan–Meier survival curves were generated using the survival (v3.6.4), survminer (v0.4.9), and survplot (v0.0.7) R packages. Statistical significance of survival differences was determined using the Fleming– Harrington (FH) test.

## Data availability

The datasets used in this study are all publicly available. The HG008 tumor/normal paired cell line is part of the Cancer Genome in a Bottle (CGIAB) project, with benchmark data and sequencing results accessible through the NIST website (https://www.nist.gov/programs-projects/cancer-genome-bottle). The COLO829 melanoma tumor/normal dataset includes PacBio HiFi data available from the PacBio portal (https://www.pacb.com/connect/datasets/) and ONT R10.4.1 data accessible via EPI2ME (https://labs.epi2me.io/colo-2024.03/). The HCC1395, HCC1954, and H2009 tumor/normal cell lines are part of the CASTLE (Cancer Standards Long-read Evaluation) project, with raw sequencing data available under BioProject accession PRJNA1086849 and related information described at https://github.com/CASTLE-Panel/castle. Additional transcriptomic datasets were obtained from The Cancer Genome Atlas (TCGA) for lung adenocarcinoma via the UCSC Xena data portal (https://xenabrowser.net/datapages/), and microarray datasets GSE40791, GSE43458, and GSE31210 were downloaded from the Gene Expression Omnibus (GEO).

## Code availability

Visit the SomaSV (v.0.0.1) repository at https://github.com/eioyuou/SomaSV to download the tool and access its documentation.

## Acknowledgements

This work has been partially supported by the National Natural Science Foundation of China (62225109, 62472120), the National Key R&D Program of China (2024YFC3406303).

## Author contributions

G.W. and T.J. designed and supervised the research. R.G. developed the SomaSV algorithm and performed the performance evaluation. H.H. and M.Z contributed to the assessment and analysis of the framework. Y.Z. and S.C. contributed to the sequencing data processing. R.G. and T.J. wrote the paper with input from all other authors. All authors read and approved the final manuscript.

